# Bone marrow stromal cells from a GDF5 origin produce zonal tendon-to-bone attachments following anterior cruciate ligament reconstruction

**DOI:** 10.1101/532002

**Authors:** Yusuke Hagiwara, Felix Dyrna, Andrew F Kuntz, Douglas J Adams, Nathaniel A Dyment

## Abstract

Following anterior cruciate ligament (ACL) injury, the ligament is often reconstructed with a tendon graft passed through bone marrow tunnels. This procedure results in a staged repair response where cell death occurs in the tendon graft, the graft is repopulated by host cells outside the graft, and tendon-to-bone attachments form at the graft/bone interface. While this healing process is well appreciated, the biological mechanisms that regulate it including the cellular origin of the repair response is poorly understood. Embryonic progenitor cells expressing growth and differentiation factor 5 (GDF5) give rise to several mesenchymal tissues in the joint and epiphyses. Therefore, we hypothesized that cells from a GDF5 origin, even in the adult tissue, would give rise to cells that contribute to the stages of repair following ACL reconstruction. ACLs were reconstructed in Gdf5Cre;R26R-tdTomato lineage tracing mice to monitor the contribution of Gdf5Cre;tdTom+ cells to graft revitalization and examine the extent to which these cells are capable of creating mineralized attachments within the bone tunnels. Anterior-posterior drawer tests were used to establish the stability of the knee following the procedure and demonstrated 58% restoration in anterior-posterior stability. Following reconstruction, Gdf5Cre;tdTom+ cells within the bone marrow expanded by 135-fold compared to intact controls in response to the injury. These cells migrated to the tendon graft interface, repopulated regions of the graft, and initiated tendon-to-bone attachments. These cells continued to organize and mature the attachments yielding a zonal insertion site at 4 weeks with collagen fibers spanning across unmineralized and mineralized fibrocartilage and anchored to adjacent bone. The zonal attachment possessed organized tidemarks with concentrated alkaline phosphatase activity similar to normal tendon or ligament entheses. This study established that mesenchymal cells from a GDF5 origin contribute to the creation of zonal tendon-to-bone attachments within bone tunnels following ACL reconstruction. Future studies will target this cell population to modulate the repair response in order to better understand key biological mechanisms that regulate tendon-to-bone repair.

## INTRODUCTION

Approximately 30% of U.S. adults suffer from tendon and ligament injuries^1^. These injuries frequently occur near insertion sites into bone (i.e., entheses) and do not spontaneously heal^2,3^. Traditional tendon-to-bone repair, where the tendon is reattached to underlying bone with sutures, does not create a zonal enthesis with collagen fibers spanning across unmineralized and mineralized fibrocartilage. On the other hand, ligament reconstructions often utilize a tendon graft placed though bone tunnels, which does produce zonal attachments. Therefore, a better understanding of the mechanisms that drive zonal tendon-to-bone attachments following ligament reconstructions may be informative to more severe tendon enthesis repairs such as to the rotator cuff tendons.

Anterior cruciate ligament (ACL) reconstruction is the most common ligament reconstruction procedure, accounting for approximately 250,000 annually in the United States alone ^4^. The annual costs of MRI, surgery, bracing and rehabilitation are estimated to exceed 2 billion dollars. Additionally, indirect costs such as lost wages, decreased productivity, and disability are substantial^4^. The highest injury prevalence is in young adult patients, typically athletes. Return to sport following ACL injury, reconstruction, and rehabilitation often exceeds eight months^5^. Therefore, there is a clinical desire to expedite this process through novel treatment strategies.

Three stages of healing have been classified following ACL reconstruction^6–8^, including i) endogenous cell death leading to degeneration of the graft, ii) vascularization and revitalization of the graft when new blood vessels and fibroblasts re-populate the graft, and iii) ligamentization of the graft where the infiltrated cells remodel the extracellular matrix and take on a ligament-like phenotype. In addition, cells that revitalize the graft also anchor the graft to bone adjacent to the tunnels via zonal tendon-to-bone insertion sites. Similar to enthesis embryonic development, the desired tendon-to-bone insertion healing process will involve synthesis of collagen fibers that will need to be anchored to the underlying bone matrix, and then the cells within this matrix will need to produce unmineralized and mineralized fibrocartilage to create a zonal insertion site.

Elucidating the biological mechanisms that govern recapitulation of an enthesis can benefit from rodent models. For instance, rat ACL reconstruction models using non-GFP allografts into GFP recipients demonstrated clearly that cell populations from outside the graft drive tunnel integration. In addition, a key signaling pathway in enthesis embryogeneis, hedgehog signaling, was shown to be active in tunnel integration in a rat ACL reconstruction model ^9^. ACL reconstruction in mice has been established ^10–12^, which opens up the possibility of using powerful genetic tools found in various mouse strains to better define the molecular regulation of tendon-to-bone repair.

Growth and differentiation factor 5 (GDF5) is a cell signaling ligand that is part of the TGFβ superfamily. It is a key signaling molecule in the formation of joints ^13^ as it is one of the earliest markers delineating the interzone cells that give rise to tissues in the joint from adjacent cartilage cells. In addition, it regulates secondary ossification and is expressed by mesenchymal cells within the secondary ossification center{Pregizer:2018el}. These cells persist into adulthood, giving rise to trabecular bone within the epiphyses^14^. Since mesenchymal cells within the joint and epiphysis originate from a GDF5 origin, this marker could be used to trace cell populations that contribute to ACL graft repopulation and tunnel integration. Therefore, the objective of this study was to Gdf5Cre transgenic mice to trace the origin of cells that revitalize the tendon graft following ACL reconstruction in a mouse model and to examine the extent to which these cells are capable of creating mineralized attachments within the bone tunnels.

## MATERIALS AND METHODS

### Mouse Models

#### GDF5Cre;R26R-tdTomato

Constitutive GDF5Cre mice^14,15^ were crossed with Ai9 R26R-tdTomato Cre reporter mice (B6;129S6-Gt(ROSA)26Sor^tm9(CAG-tdTomato)Hze^/J, Jackson Labs) to label mesenchymal cells within the joints and epiphyses to monitor revitalization of ACL grafts following reconstruction.

### Experimental Design

Three to four month old mice were used in this study. Three groups were created for anterior-posterior drawer mechanical assessment to establish the stability of the knee in this new model: ACL reconstructed knee (n=8), ACL-deficient knee (transected ACL: ACLT) (n=9), or intact knee (n=9). The animals were assessed at 4-weeks post-surgery. Additional mice were subjected to ACL reconstruction and the bone tunnel regions were examined using multiplexed mineralized cryohistology at 1-week (n=3) and 4-weeks (n=4) post-surgery. These mice received an intraperitoneal (IP) injection of demeclocycline (30μg/g) one day prior to sacrifice to label deposited mineral.

### ACL Reconstruction Procedure

All mice were approved by UConn Health IACUC committee. Mice were anesthetized and sterilely prepped. Three small incisions (3mm each, spaced ~20mm apart) were made in the tail to access the tail tendons (Fig. 1A). Continuous tail tendon bundles (3-4 tendons in each bundle) were then identified and removed (3-4cm in length)(Fig. 1B). Suture was passed through a small length of 25G needle (3mm long) to create a cortical fixator which was then passed over the tail tendon bundle (yellow arrowhead in Fig. 1C). Nylon suture was tied around the tendons at each end of the bundle (green circle in Fig. 1C) and then the bundle was submerged in PBS until the tunnels were drilled. The knee joint space was accessed via anteromedial approach (Fig. 1D) and the ACL was transected near its femoral insertion with a 28G needle. A manual drawer test was performed to confirm that the ACL was cut while not damaging the PCL. The tibial tunnel was drilled retrograde from the ACL tibial insertion footprint to the outside of the medial tibia with a 28G needle. The needle was removed and then reinserted antegrade through the tibial tunnel to drill the femoral tunnel retrograde from the femoral ACL insertion footprint (Fig. 1E-F). The tendon graft was fed through the femoral and tibial tunnels, respectively, such that the cortical fixator was positioned on the outer cortex of the femur (Fig. 1G-H). At the tibial tunnel exit, the two ends of the tail tendon bundles were tied together with a basic square knot and then sutured to adjacent muscle (Fig. 1I). Another manual drawer test was performed to confirm that anterior-posterior translation was stabilized. Finally, the incision was closed with suture, analgesia was initiated (0.1 mg/kg buprenorphine), and the animal was returned to its cage following recovery from anesthesia.

**Fig. 1:**
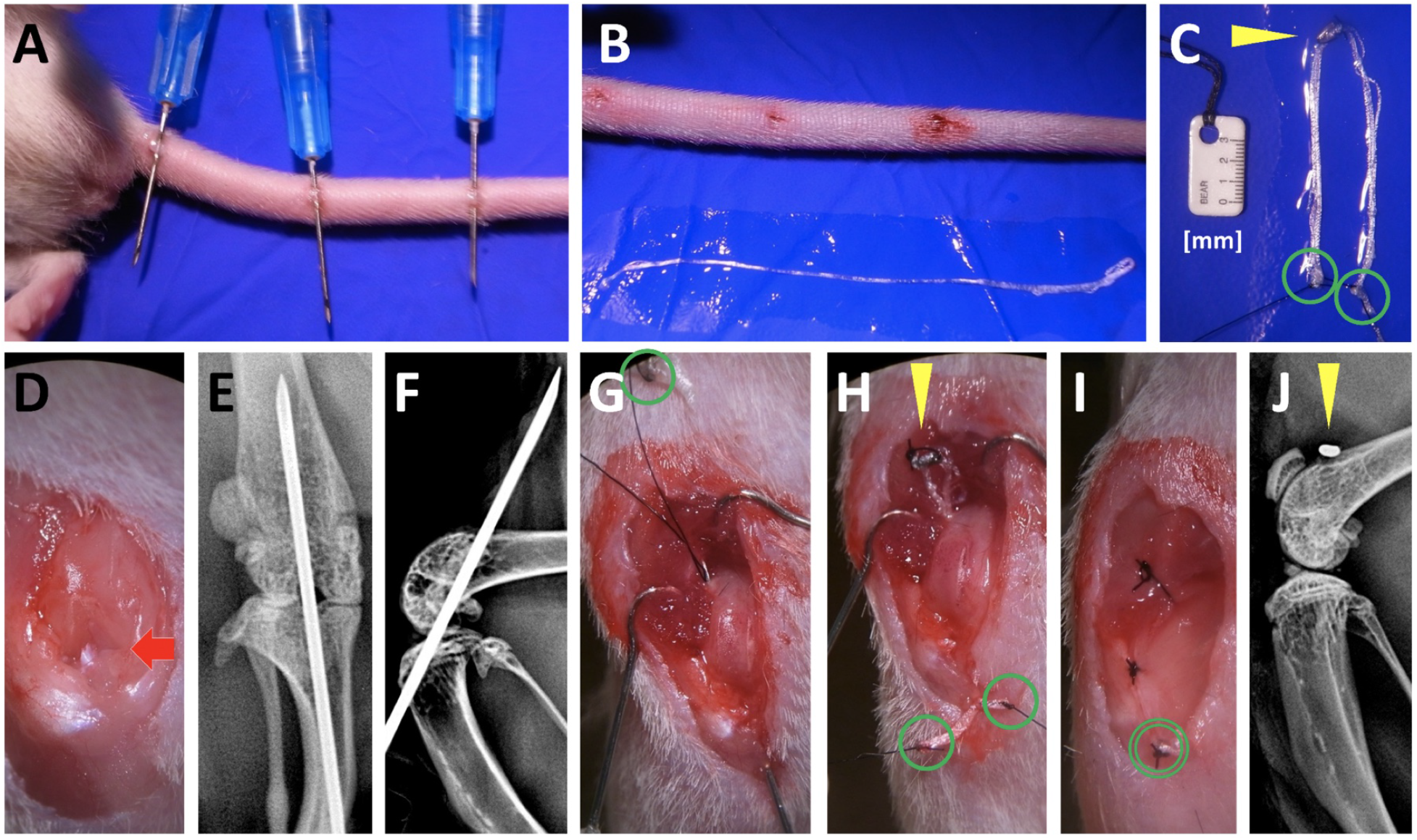
Murine ACL reconstruction procedure using tail tendon autograft. Three incisions along the length of the tail (A) were used to isolate 3-4cm segments of tail tendons (B). These tendon bundles were secured with suture (green circles in C) and a cortical fixator was applied to the midpoint of the bundle (yellow arrowhead in C). The ACL (red arrow in D) was accessed using an anteromedial approach. The tibial and femoral tunnels were drilled (E-F) and the graft was fed into the tunnels from the femur (G) and out the tibia (H). The cortical fixator (yellow arrowhead in H and J) provided fixation on the femur while the tail tendon bundle was tied and sutured to the adjacent muscle (I) to provide fixation on the tibia.

### Anterior-Posterior Drawer Test

Both hindlimbs were harvested at four weeks post-surgery. The tibia was pinned to a small, rigid foam block by passing 25-27G needles transversely through the bone. Needles were then passed through the femur in a similar fashion. The tibia block was taped down to the horizontal shelf within a digital X-ray cabinet (Faxitron LX-60) and the knee was positioned at 90° of flexion. Sutures were wrapped around the needles in the femur then attached to a 10g weight that was hung off the edge of the Faxitron shelf via a pulley. The 10g force was applied in the tibial anterior direction and an X-ray image was acquired to define the posterior limit of motion. Subsequently, the weight was applied in the posterior direction of the tibia and an additional X-ray image was acquired to define the anterior limit of A-P drawer. The images were assembled as layers in image analysis software. The distance between the anterior surface of the femoral condyle in the posteriorally- and anteriorally-displaced images was measured and normalized by the width of the tibial plateau in the sagittal plane to quantify anterior-posterior drawer. Cryohistology Hindlimbs were fixed in 10% neutral buffered formalin for 1-2 days, transferred to 30% sucrose in PBS overnight, and embedded in OCT. The knee was sectioned in the sagittal plane to capture longitudinal sections of the tibial and femoral tunnels. All sections were made from undecalcified joints using cryofilm 2C^14,16–20^, which maintains morphology of mineralized sections. The taped sections were secured to glass microscope slides using chitosan adhesive and rehydrated with 1X PBS prior to imaging.

### Alkaline phosphatase staining

Alkaline phosphatase (AP) staining was performed using the Vector blue alkaline phosphatase substrate kit (Vector Labs) according to manufacturer protocols. The sections were incubated in the substrate solution for 15 minutes. The AP signal was imaged using a Cy5 filter (Ex:640/30, Em:690/50).

### Imaging

Three rounds of imaging were performed on each section. This sequence was possible because the cryofilm tape adheres to the tissue and allows for the coverslip to be removed between imaging steps without damaging the section. The order of imaging included: (1) fluorescent reporters and fluorescent mineralization labels, (2) alkaline phosphatase staining with DAPI counterstain, and (3) toluidine blue O (0.025% in dH2O) staining.

### Image analysis to quantify expansion of Gdf5Cre;tdTom+ cells

To measure the extent to which the Gdf5Cre;tdTom+ cells within the intact marrow expanded in response to the tunnel injury, the percent area of tdTom+ signal was quantified from tissue sections by drawing a region of interest that included all of the tibial epiphyseal bone marrow. An equivalent minimum threshold value was set for all samples and the positive pixels were divided by total pixels to calculate percent area.

### Statistics

Anterior-posterior drawer measurements were compared via Mann Whitney U tests with significance level set to p < 0.017 to account for multiple comparisons. tdTom+ percent area measurements were compared via Mann Whitney U tests with significance level set to p < 0.05.

## RESULTS

### ACL reconstruction restored 58% of the anterior-posterior stability compared to an ACL-deficient knee after four weeks of ambulation post-reconstruction

To assess knee stability following the ACL reconstruction healing process, reconstructed knees were compared to ACL-deficient (ACLT) and intact knees at four weeks post-surgery. A-P translation in intact knees corresponded to 2.9±1.6% of the tibial plateau compared to 22.8±6.7% in ACLT knees (p < 0.017). After 4 weeks post-reconstruction, A-P translation corresponded to 11.0±6.7% in the ACLR knees, which equates to 58% restoration of anterior-posterior stability compared to the ACLT group (p < 0.017).

### Gdf5Cre;tdTom+ cells are found in several tissues within the joint and epiphysis in the adult

Gdf5 is one of the key signaling factors expressed by interzone cells during joint development ^13^. Progeny of Gdf5-expressing cells are found within mature tissues in the joint even into adulthood^14^. In our samples, we similarly see Gdf5Cre;tdTom+ cells within the articular cartilage, cruciate ligaments (Fig. 3A1), collateral ligaments, synovium, and epiphysis. The cells in the epiphysis include osteoblasts overlaying a mineralization label in the trabecular bone (arrowhead in Fig. 3A2) and cells residing within the bone marrow (arrow in Fig. 3A2). The tail tendons of these mice however, contain <1% Gdf5Cre;tdTom+cells (Fig. 3B).

**Fig. 2:**
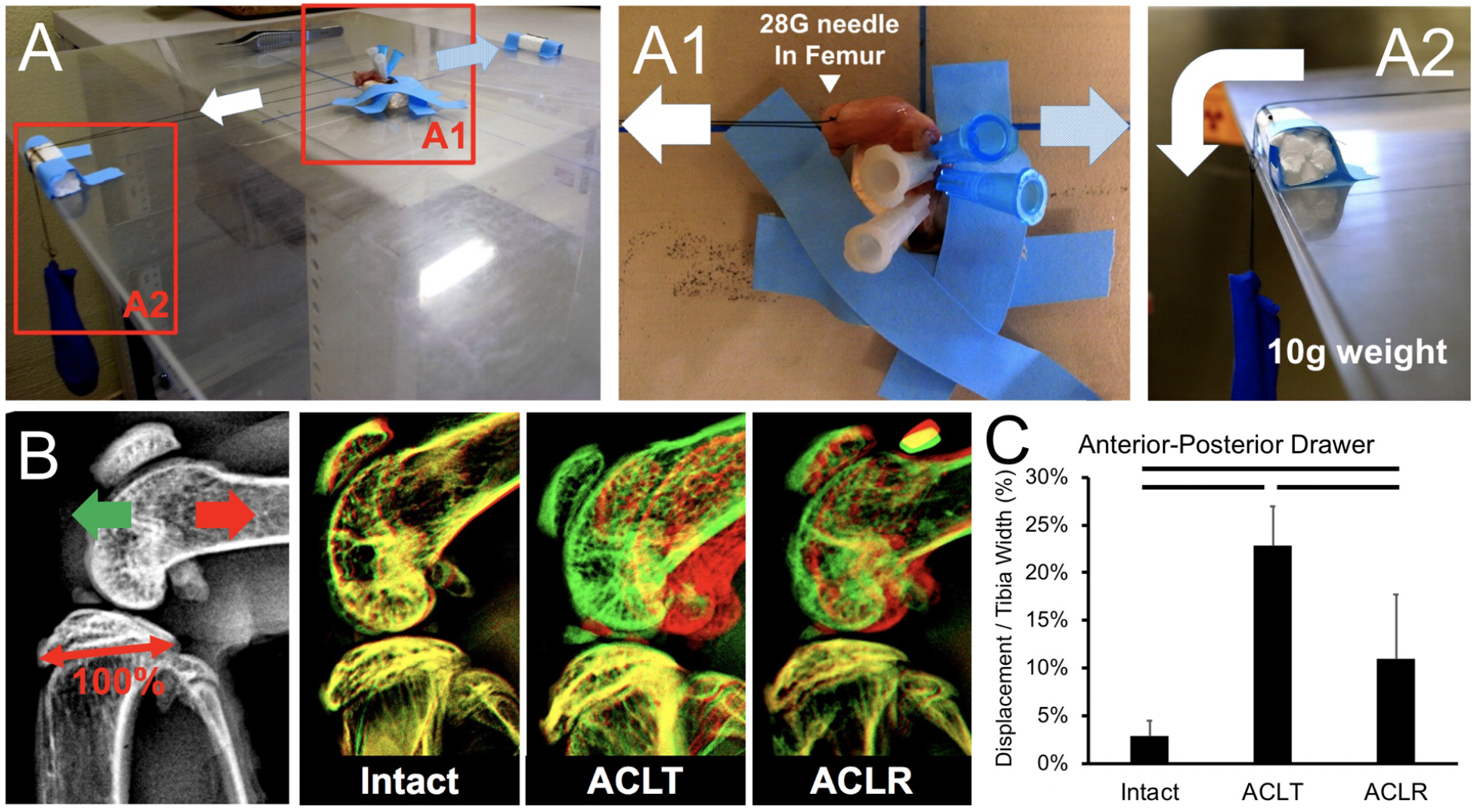
Reconstruction procedure restores 58% of anterior-posterior stability of knee joint. At 4-weeks post-surgery, limbs were harvested and the tibia was pinned to a foam block using needles and taped to the shelf of a digital X-ray cabinet (A). A 28G needle was also passed through the femur (A1) and then suture was tied to this needle on one end and to a 10g weight on the other end (A2). The weight was applied in the anterior (light blue arrow in A1 or green arrow in B) and posterior (white arrow in A1 or red arrow in B) directions relative to the tibia, acquiring a radiograph at each position. The two images were overlaid in image analysis software (B) and the displacement of the femur from anterior to posterior images was measured and normalized to the width of the tibial plateau (denoted as 100% in B). Measurements are reported in (C). Bars indicate p < 0.017.

**Fig. 3:**
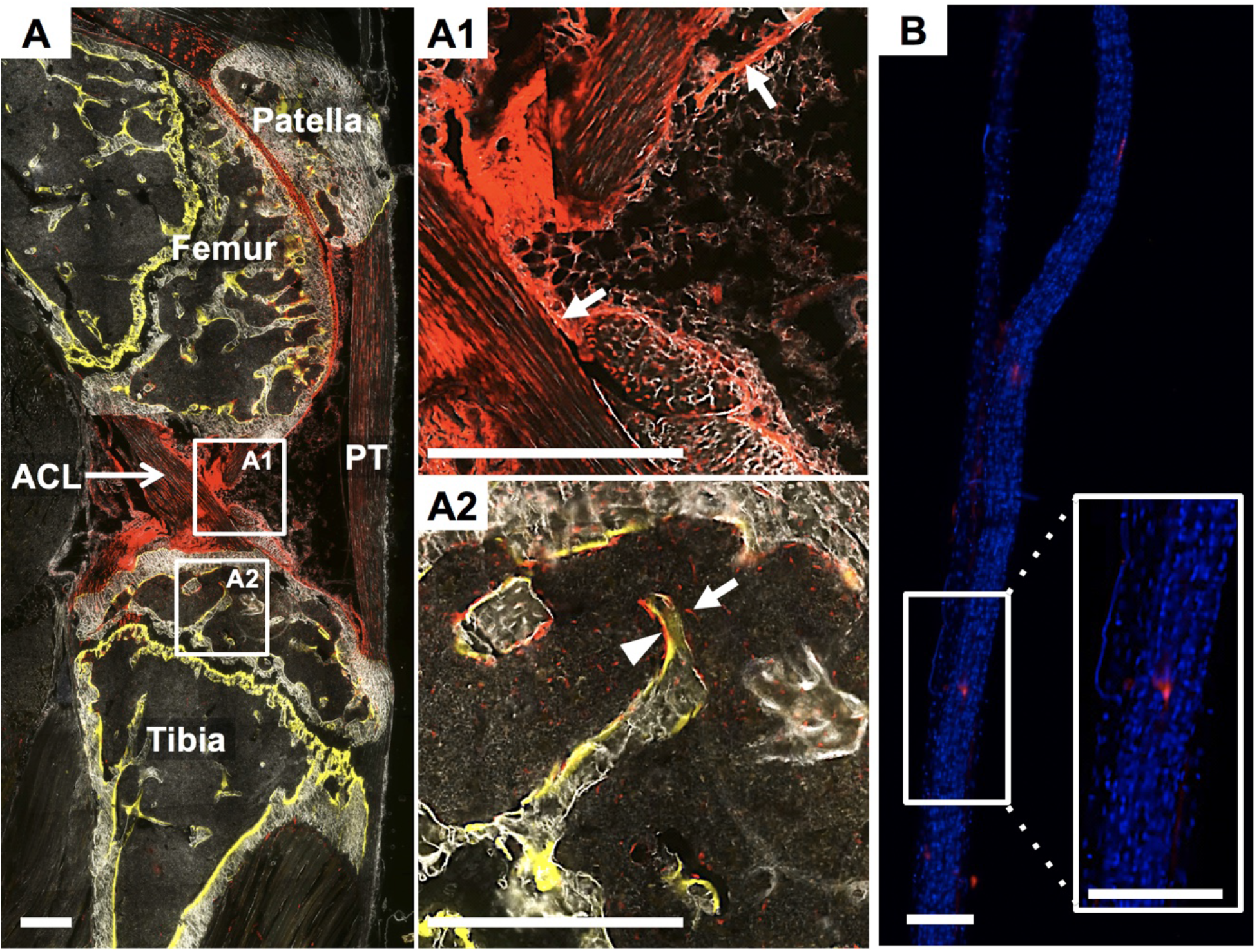
tdTom+ cells exist within the knee but not tail tendons of Gdf5Cre;R26R-tdTomato mice. The intact contralateral knee from a ACLR mouse that received demeclocycline injection (yellow in A) the day prior to sacrifice. Gdf5Cre;tdTom+cells exist within the synovial lining (white arrows in A1), cruciate ligaments (A1), epiphyseal bone (arrow head in A2), and bone marrow (arrow in A2). Conversely, the tail tendons from these mice possess very few Gdf5Cre;tdTom+ cells (B). Scale bars: 200μm.

### Following ACL reconstruction, Gdf5Cre;tdTom+ cells within the epiphyseal bone marrow expand in response to the injury and initiate attachments with the tendon graft

At 1 week post-surgery, there was considerable expansion of Gdf5Cre;tdTom+ cells within the bone marrow (Fig. 4A). The tdTom+ percent area within the epiphyseal bone marrow increased to 54.1 ±12.1% from only 0.4±0.3% in the intact contralateral limb (p < 0.05). The leading front of the expanding cells approached the tendon graft interface (Fig. 4A3-4) where collagen fibers via second harmonic generation (SHG) imaging (yellow arrow in Fig. 4A5) containing tdTom+ cells were juxtaposed with the denser and more aligned collagen fibers of the tendon graft.

**Fig. 4:**
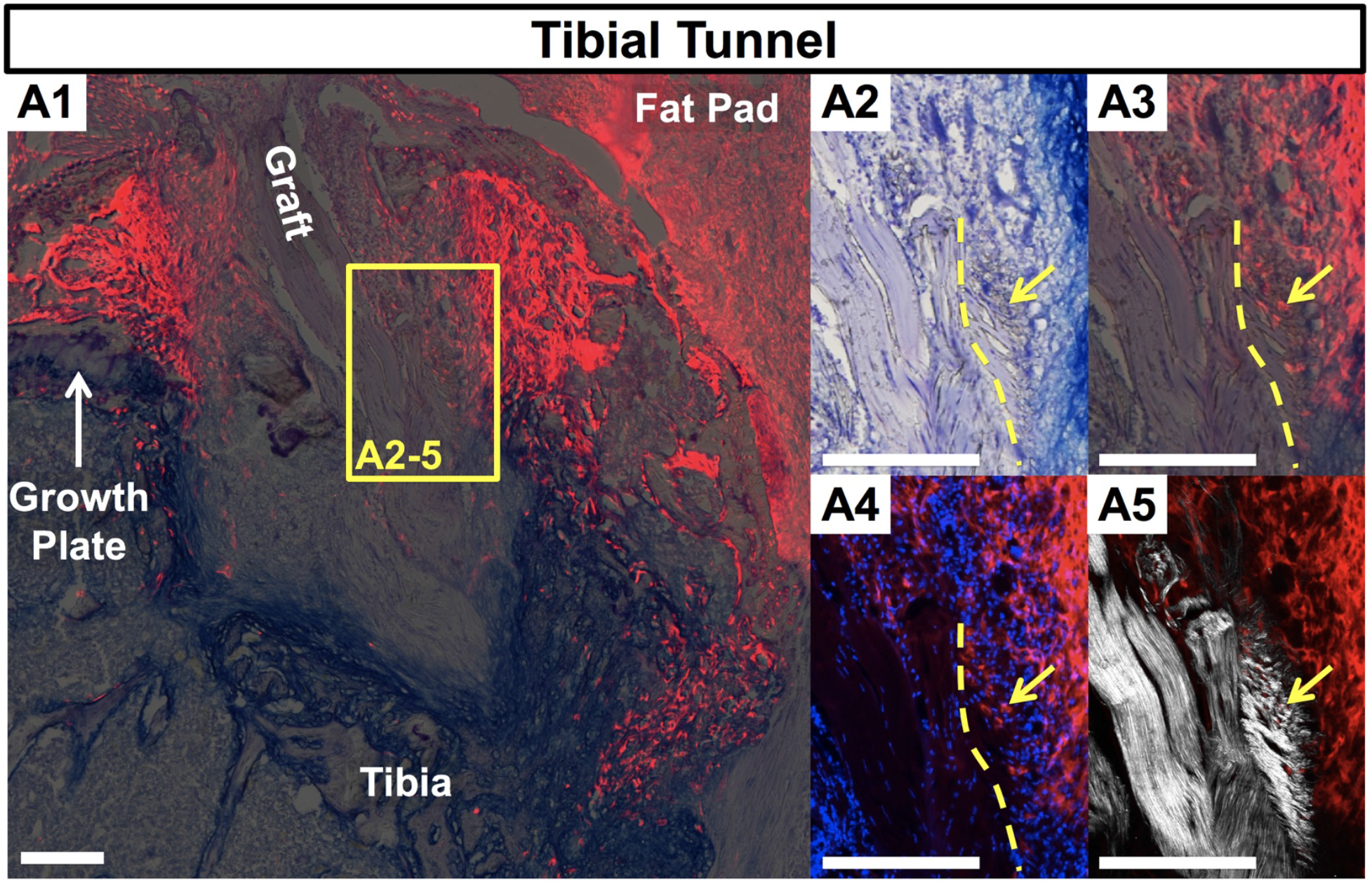
Expansion of Gdf5Cre;tdTom+ epiphyseal bone marrow cells and early attachments at 1-week post-surgery. A) Tibial tunnel with tendon graft in the center flanked by expanding tdTom+ bone marrow population. Panels A2-5 demonstrate early attachments at the interface of the graft with the marrow space. A2) toluidine blue O stain, A3) tdTom channel (red) overlaid onto toluidine blue image, A4) tdTom channel with DAPI counterstain, and A5) collagen SHG with tdTom channel. Dotted lines denote interface between dense collagen of graft and looser collagen containing tdTom+ cells (yellow arrow).

### Gdf5Cre;tdTom+ cells give rise to mineralized attachments within the bone tunnels

The Gdf5Cre;tdTom+ bone marrow cells that interfaced with the tendon graft at 1 week post-surgery were now embedded between dense collagen fibers spanning an organized tidemark (yellow demeclocycline label denoted by white arrow in Fig. 5A3-6) separating unmineralized and mineralized fibrocartilage (* in Fig. 5A3) of zonal attachments. Alkaline phosphatase facilitates the deposition of mineral and is concentrated near the tidemark in the normal enthesis ^17^. Alkaline phosphatase was concentrated in the bone tunnel attachments as well (Fig. 5A6) and was found surrounding stacked fibrochondrocytes of the unmineralized fibrocartilage adjacent to the tidemark (Fig. S1B4). The collagen fibers of the attachments were anchored to the underlying bone which surrounded the tunnel at 4 weeks. In contrast, very little trabecular bone was found at the graft interface at 1 week post-surgery. Attachments with all of these attributes were found within both tunnels for all mice at four weeks post-reconstruction.

**Fig. 5:**
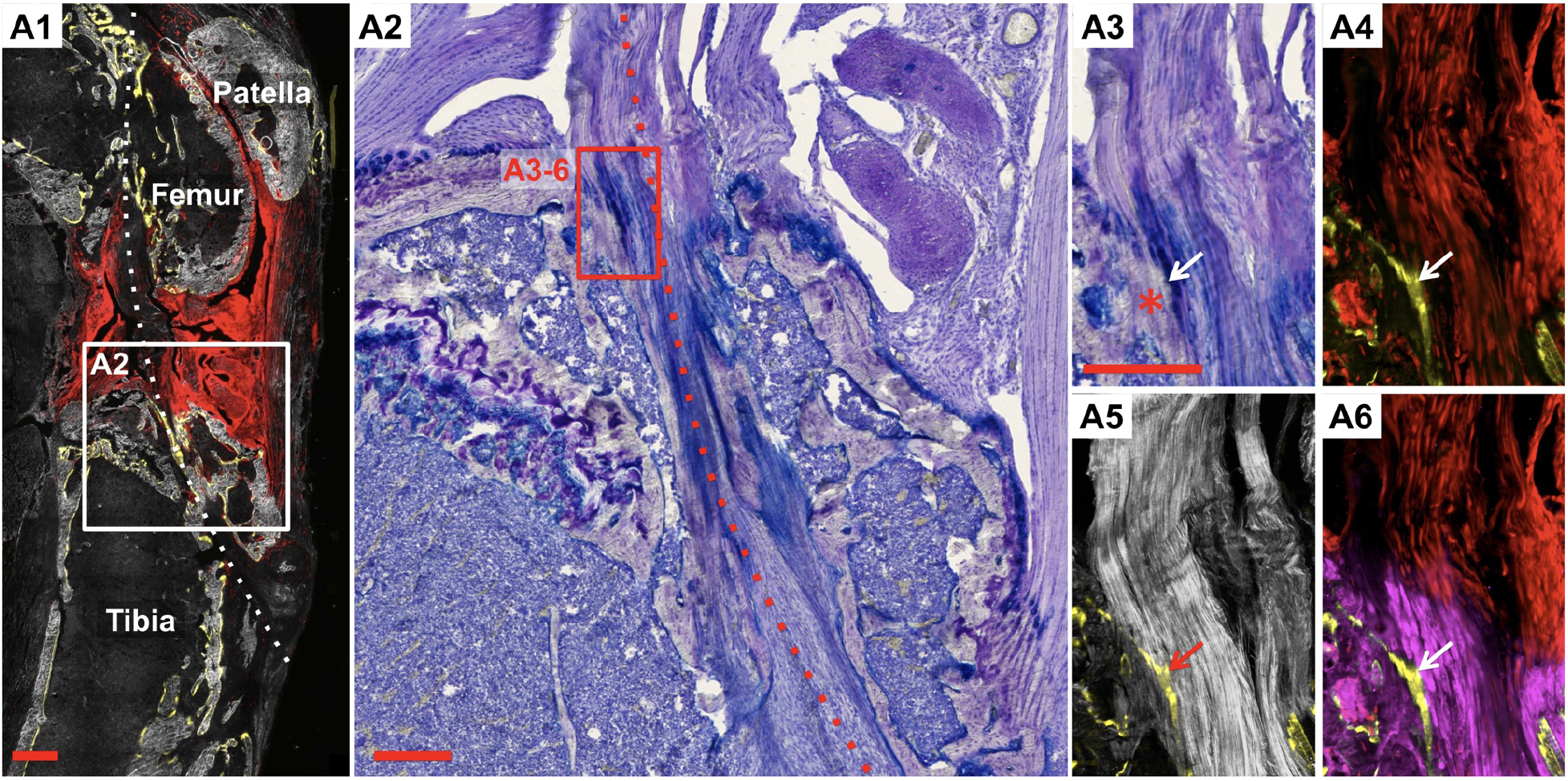
Gdf5Cre;tdTom+ cells produce mineralized attachments in the bone tunnel. A1) Image of entire knee joint with dotted line denoting tunnel path at 4 weeks post-surgery. A2) Toluidine blue stained image of tibial tunnel. Panels A3-6 demonstrate zonal attachment at entrance to tibial tunnel. A3) Toluidine blue image, A4) tdTom channel (red) with demeclocycline label (yellow), A5) collagen SHG with demeclocycline, and A6) tdTom channel with demeclocycline and alkaline phosphatase activity (magenta). Arrows denote yellow demeclocycline tidemark. * denote mineralized fibrocartilage. Scale bar: 200μm.

### Gdf5Cre;tdTom+ cells revitalized the tendon graft within the bone tunnels prior to the graft midsubstance within the joint space

At one week following reconstruction, the tendon graft was hypocellular both within the bone tunnels (Fig. 4A1,4) and the joint space (Fig. 6A), with the majority of nuclei in the graft existing near the graft surface (white arrow). By 4 weeks, a large number of Gdf5Cre;tdTom+ cells were found within the tendon graft, especially in the regions of the graft within the bone tunnels (Fig. 4A4 and S1B3). In contrast, the graft tissue within the central region of the joint remained largely acellular at 4 weeks (Fig. 6B). The majority of Gdf5Cre;tdTom+ cells that infiltrated the graft within the joint space were adjacent to the tunnel entrances on both the femur and tibia (yellow arrow in Fig. 6B).

**Fig. 6:**
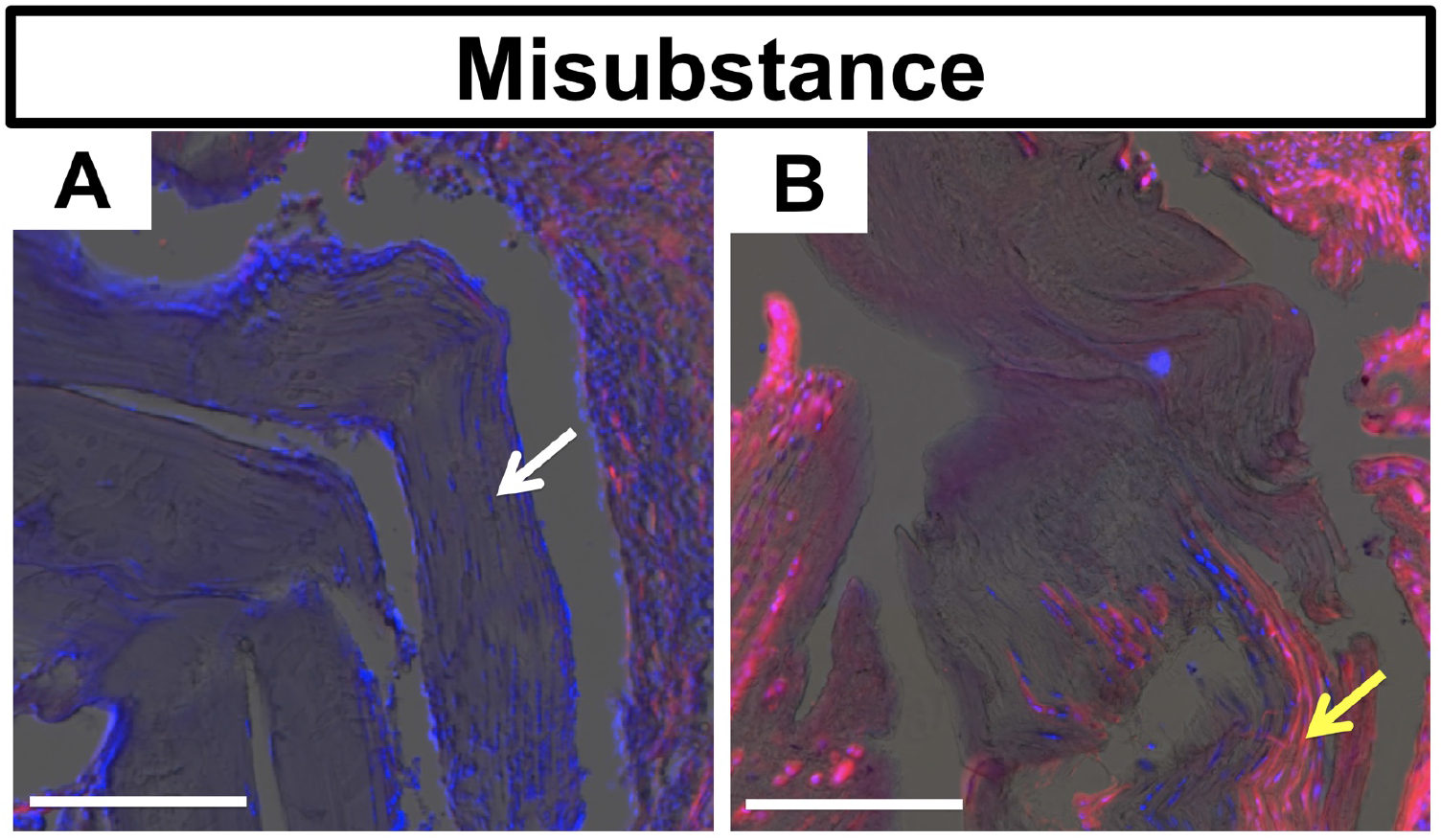
Cell death within graft midsubstance and limited cell repopulation. Images of tendon graft within joint space at 1-week (A) and 4-weeks (B) post-surgery. Images are of tdTom channel (red) with DAPI counterstain (blue) overlaid on toluidine blue image. Graft at 1 week possesses live cells near the surface of the graft (white arrow), while the graft was mostly acellular at 4 weeks with cellular portions coming from repopulating Gdf5Cre;tdTom+ cells (yellow arrow). Scale bars: 200μm.

## DISCUSSION

Two events are crucial to successful biological repair following ACL reconstruction: 1) cells need to revitalize the graft, as resident cells within the graft frequently die following reconstruction, and 2) the soft tissue graft needs to integrate with the adjacent bone in the tunnels through the formation of zonal attachments^6,21^. We developed this mouse ACL reconstruction model, which is similar to previous murine ACLR models ^10–12^, to monitor graft revitalization following reconstruction and the formation of mineralized attachments within the bone tunnel (Figs. 4-6). We performed these procedures in Gdf5Cre;R26R-tdTomato mice because progeny of Gdf5-expressing embryonic cells give rise to mesenchymal cells within the joint and epiphysis but not the tail tendons, permitting us to monitor infiltration of surrounding resident cells into the unlabeled tail tendon (Fig. 3B). We found in this model that cells from outside the graft within the bone marrow are the primary cell population to revitalize the graft within the bone tunnels, while cells within the synovial lining revitalize the graft midsubstance within the joint space. In addition, these cells from a Gdf5 origin have the potential to form zonal tendon-to-bone attachments within the bone tunnels.

The coordinated events of zonal enthesis formation are well characterized in the developing animal during embryogenesis and postnatal growth. Enthesis progenitors expressing Gdf5^14,15,17^, among other markers^22–24^, go on to assemble collagen fibers that are anchored to the underlying cartilage template, surround these fibers with a proteoglycan-rich matrix (i.e., umineralized fibrocartilage), and then finally mineralize a portion of this fibrocartilage to create the zonal enthesis. In this study, we found that adult cells from a Gdf5 origin within the epiphyseal bone marrow had the potential to assemble zonal attachments within the bone tunnel following ACL reconstruction. While not as organized, these attachments displayed aspects of a normal zonal enthesis, with collagen fibers spanning across an organized tidemark (Fig. 5A5), alkaline phosphatase activity adjacent to this tidemark (Fig. 5A6), and proteoglycans within the fibrocartilage (purple stain in Fig. 5A3). Therefore, this murine ACL reconstruction model could be used to study specific molecular mechanisms that govern adult zonal tendon-to-bone repair.

There were key aspects of the zonal attachment formation in the ACL reconstruction model that are quite distinct from normal development. Even though the adult bone marrow mesenchymal progenitor cells that give rise to these attachments come from a Gdf5 lineage, they are quite different from the interzone-derived cells that give rise to the normal ACL enthesis. The Gdf5Cre;tdTom+ cells that activate in response to the bone tunnel injury give rise to both the mineralized attachments in the tunnel as well as the contiguous bone adjacent to the attachment (Fig. S1B2). Therefore, there is either a fate decision that is made by a common progenitor, or there is a heterogenous progenitor pool with some cells serving as osteoprogenitors giving rise to trabecular bone and others as chondroprogenitors giving rise to the enthesis. Defining that a heterogenous progenitor pool exists versus identifying the mechanisms that drive fate decisions from a common progenitor would provide novel cell and molecular targets to enhance or accelerate the repair response.

The mechanisms that drive maturation of the enthesis during postnatal growth may be similar to the maturation process that occurs in the tunnel attachments, especially since these attachments have mineralized fibrocartilage with an organized tidemark. The hedgehog pathway is one of the key signaling pathways that drives maturation and mineralization of the enthesis during development^17,25–29^, as ablation of this pathway resulted in severe reductions in mineralized fibrocartilage formation. Indian hedgehog (IHH) is the ligand expressed during mineralization of the enthesis ^17^, more so than sonic or desert hedgehog, so this isoform likely regulates the maturation process. Recent studies in rat and murine ACL reconstruction models demonstrated that key hedgehog signaling factors including IHH, Ptch1, and Gli1 were expressed in the mineralized attachments in the bone tunnels^9,12^. However, it is not known if hedgehog serves a similar function in promoting mineralized fibrocartilage formation in these adult attachments as it does in the developing enthesis because genetic ablation studies have not been performed. Murine models of ACL reconstruction, however, provide a platform to modulate the pathway in a similar fashion to the pivotal enthesis developmental studies. Future studies will work to modulate the GDF5-origin cells that contribute to tunnel integration to better understand the molecular mechanisms driving zonal tendon-to-bone repair.

## ACKNOWLEDGEMENTS

The authors would like to thank Dr. David Kingsley for graciously providing the GDF5Cre mice. This work was supported by NIH grant K99/R00 AR067283 (NAD) and DOD grant W81XWH-15-1-0371 (DJA)

## COMPETING INTERESTS

The authors declare no competing financial interests.

## SUPPLEMENTAL FIGURE

**Fig. S1:**
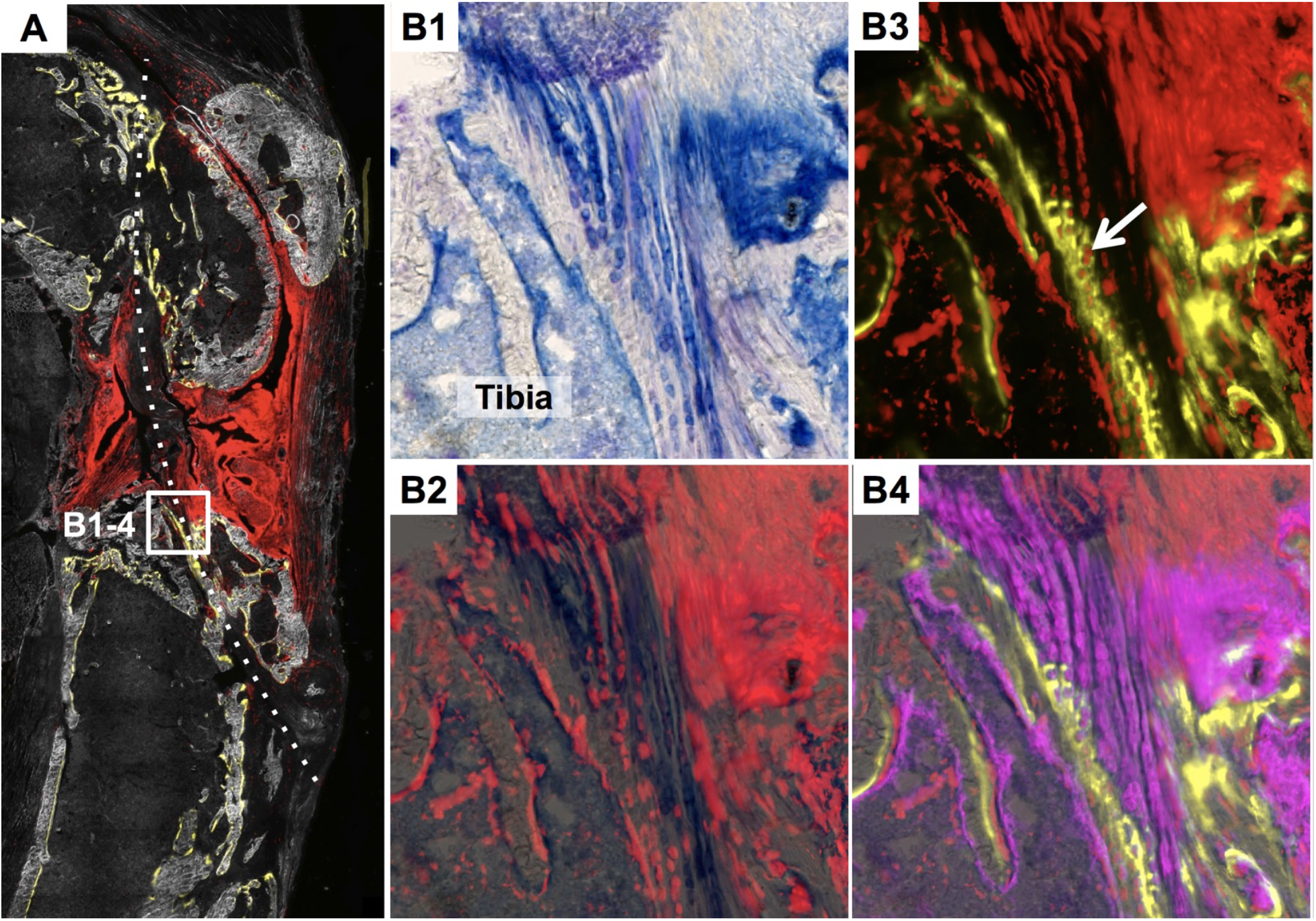
Additional example of mineralized attachment at 4 weeks with stacked fibrochondrocytes. A) Overview image of entire knee with dotted line denoting tunnel path. B1) Toluidine blue image of attachment in tibial tunnel, B2) tdTom (red) overlaid onto toluidine blue image, B3) tdTom (red) with demeclocycline label (yellow), B4) tdTom (red) with demeclocycline label (yellow) and alkaline phosphatase (magenta) overlaid on toluidine blue image.

